# GNOMES: an integrated framework for genome-wide normalization and differential binding analysis of CUT&RUN and ChIP-seq data

**DOI:** 10.64898/2026.04.16.718722

**Authors:** Thomas Roule, Naiara Akizu

## Abstract

**Background:** Despite their use, quantitative comparison of epigenomic datasets such as ChIP-seq and CUT&RUN remains challenging, particularly due to difficulties in signal normalization across samples and conditions. Normalization solely based on sequencing depth is often insufficient due to the high variability in signal-to-noise ratios across samples, even from a same experiment. While exogeneous spike-in normalization can address some issues, robust spike-in controls are not always available, and may introduce additional experimental burden and computational complexity. Furthermore, normalization and differential binding analysis are typically performed using separate bioinformatics tools. Indeed, most differential analysis frameworks operate on raw count matrices, preventing users from visually inspecting normalized signal tracks and evaluating how normalization influences the results. To overcome these challenges, we developed *GNOMES (Genome-wide NOrmalization of Mapped Epigenomic Signals)*, a framework that integrates signal normalization, quality control, and differential binding analysis within a unified workflow.

**Results:** *GNOMES* is a user-friendly tool able to process ChIP-seq and CUT&RUN datasets from aligned reads, and generate normalized coverage profiles and differential binding results. The tool implements a robust genome-wide normalization strategy based on percentile scaling of signal local maxima, enabling stable normalization between biological replicates and conditions. *GNOMES* supports both single- and paired- end sequencing, does not required a negative control (input or IGG), and can be applied to both broad (histone marks) or narrow (transcription factor) enrichment patterns. The workflow includes normalization, optional consensus peak identification, and differential binding analysis. For each step, *GNOMES* generates extensive quality-control metrics and visual outputs, including normalized bigWig tracks, median signal tracks, BED files of regions with significant changes, and diagnostic plots such as heatmaps and PCA. *GNOMES* is highly configurable and integrates established tools such as MACS2 for candidate peak regions identification for differential binding analysis, as well as DESeq2 and edgeR for statistical testing. Finally, *GNOMES* is organism-agnostic and can be applied to epigenomic datasets from any model system.

**Conclusions:** *GNOMES* provides an integrated and highly customizable environment for normalization and differential binding analysis of epigenomic sequencing data. By integrating signal normalization, with downstream differential statistical method for differential binding analysis, and comprehensive quality control, *GNOMES* simplifies the analysis of ChIP-seq and CUT&RUN datasets, for the identification of chromatin changes.

## Background

Chromatin profiling techniques such as ChIP-seq and CUT&RUN are widely used to map binding of transcription factors, and histone marks genome-wide [1]. These approaches enable the identification of genomic regions enriched for specific chromatin features, providing insight into chromatin structure and gene regulation. These techniques can also be applied in multiple biological conditions, such as wild-type versus mutant samples or untreated versus treated cells, in order to identify regions with different chromatin features between conditions. However, accurate quantitative comparison of mapped epigenomic signals across conditions remains challenging. Variability in sequencing depth and poor signal-to-noise ratio often introduce substantial differences in signal intensity between samples/replicates, even from the same experiments. Thus, robust normalization of signal is a critical step for downstream differential binding analysis [2].

Two main, interdependent, strategies are commonly used to identify differential binding regions from epigenomic datasets. The first relies on genome-wide sliding window approaches, in which the entire genome is scanned using fixed bins and statistical testing is performed on read counts within each bin. Tools such as rgt-THOR and csaw, use this strategy to identify regions showing significant changes in signal between conditions [3, 4]. While sliding window -based approaches allow unbiased genome-wide detection of changes, they can be sensitive to background noise and may identify significant changes in regions with low-signal areas. As the entire genome is tested, small differences between near-zero signals can sometimes appear statistically significant, notably in datasets with poor signal-to-noise ratios [5]. The second strategy follows a peak-centric approach, in which enriched regions, (i.e. peaks), are first identified using algorithms such as MACS2, or SICER [6, 7], and read counts within these peaks are quantified and compared between conditions by applying statistical methods such as DESeq2 or edgeR [8, 9]. This approach focuses on biologically meaningful regions (e.g., true peaks), and is therefore more robust for datasets with substantial background noise [5]. Despite the availability of these methods, current workflows often separate signal normalization, peak identification, and differential binding analysis into independent steps, frequently requiring multiple tools and custom scripts. Furthermore, in most cases, differential testing is performed directly on the raw count matrices and normalized signal tracks suitable for visualization in genome browsers are not generated, making difficult for users to visually inspect region with significant changes, limiting the ability to validate results through visual inspection [10].

To address these limitations, we developed *GNOMES (Genome-wide NOrmalization of Mapped Epigenomic Signals)*, a flexible framework that integrate signal normalization and peak-centric differential binding analysis within a single workflow. *GNOMES* implements a normalization strategy based on percentile scaling of signal local maxima, enabling robust normalization across datasets without requiring exogeneous spike-in controls [11]. Briefly, the method rescales signal intensities so that the upper signal distribution (defined by local maxima percentiles) becomes comparable across samples, reducing differences in global signal magnitude between samples. The framework is implemented with MACS2 algorithm to identify consensus peaks [6], that serve as candidate regions for the differential binding analysis, and perform statistical testing with DESeq2 or edgeR [8, 9]. *GNOMES* is design to be broadly applicable across experimental systems. It supports ChIP-seq and CUT&RUN datasets, works with both single and paired -end sequencing, is not limited to an organism, and can be applied to both broad histone modifications, and narrow transcription factor. The tool provides extensive quality-control metrics and diagnostic visualizations, allowing users to assess data quality and normalization performance. By combining normalization, visualization, and differential analysis in a highly configurable environment, *GNOMES* aims to facilitate epigenomic datasets analysis while enabling users to tailor analytical parameters to their specific experimental features.

To illustrate *GNOMES* performance, we analyzed H3K27me3 ChIP-seq data generated from mouse cerebellum at postnatal day 12 (P12) and postnatal day 21 (P21) [12]. Differentially H3K27me3 enriched regions identified by *GNOMES* were associated with nearby genes and integrated with gene expression data to assess correlation between histone mark binding and gene expression. This integrative analysis provides a biological validation of *GNOMES* by testing whether regions showing differential enrichment correspond to expected expression changes of associated genes.

## Implementation

### System architecture

*GNOMES* provides a command-line interface designed to run on any Unix-based system. The software is primarily written in Python and integrates components implemented in Python, Bash, and R. The software can be installed using a Conda, or with Apptainer. *GNOMES* supports both Single-End (SE) and Paired-End (PE) data from ChIP-seq or CUT&RUN. *GNOMES* can be applied to any genome assembly, as the primary required inputs are aligned sequencing reads in BAM format. *GNOMES* is integrated with several widely used bioinformatics tools, including MACS2 for peak detection [6], DESeq2 and edgeR for statistical testing of differential binding [8, 9], and deepTools for signal visualization [13].

The workflow is organized into three main modules: *GNOMES norm*, *consensus*, and *diff* (**Figure1**). The norm module performs signal normalization from the aligned BAM files, and generate normalized signal tracks in BigWig format, for each sample, and also generate median signal track for each condition. The *norm* module also generates quality-control plots and metrics, including summaries of the scaling factors applied during normalization, as well as heatmap and PCA plots allowing users to assess sample homogeneity before and after *GNOMES* normalization. The consensus module is optional and allow users to identify optimal regions for differential binding analysis. It uses MACS2 to detect peaks across samples and conditions and generates multiple BED files representing different sets of consensus peaks, using different statistical threshold. These outputs enable users to visually inspect candidate regions, for example in genome browsers such as IGV, and select the most appropriate peak set for downstream differential analysis. Finally, the *diff* module performs the differential binding analysis using DESeq2 or edgeR within a genomic region provided by the user, or automatically generated. The *diff* module also generates quality-control plots and metrics, including tables of differentially bound regions with fold-change and q-value statistics, BED files of regions showing gain or loss of signal, and diagnostic PCA and heatmap plots to assess similarity of peak signals across samples. Furthermore, heatmap and signal profile plots are produced for significantly enriched regions, allowing users to visually evaluate differential binding patterns. Input control datasets (such as Input or IgG controls) are optional but can be incorporated into the workflow. During the *norm* stage, control signal can be subtracted from the immunoprecipitation (IP) signal, and during the *consensus* stage, controls can be used by MACS2 as background for peak calling [6].

**Figure 1.**
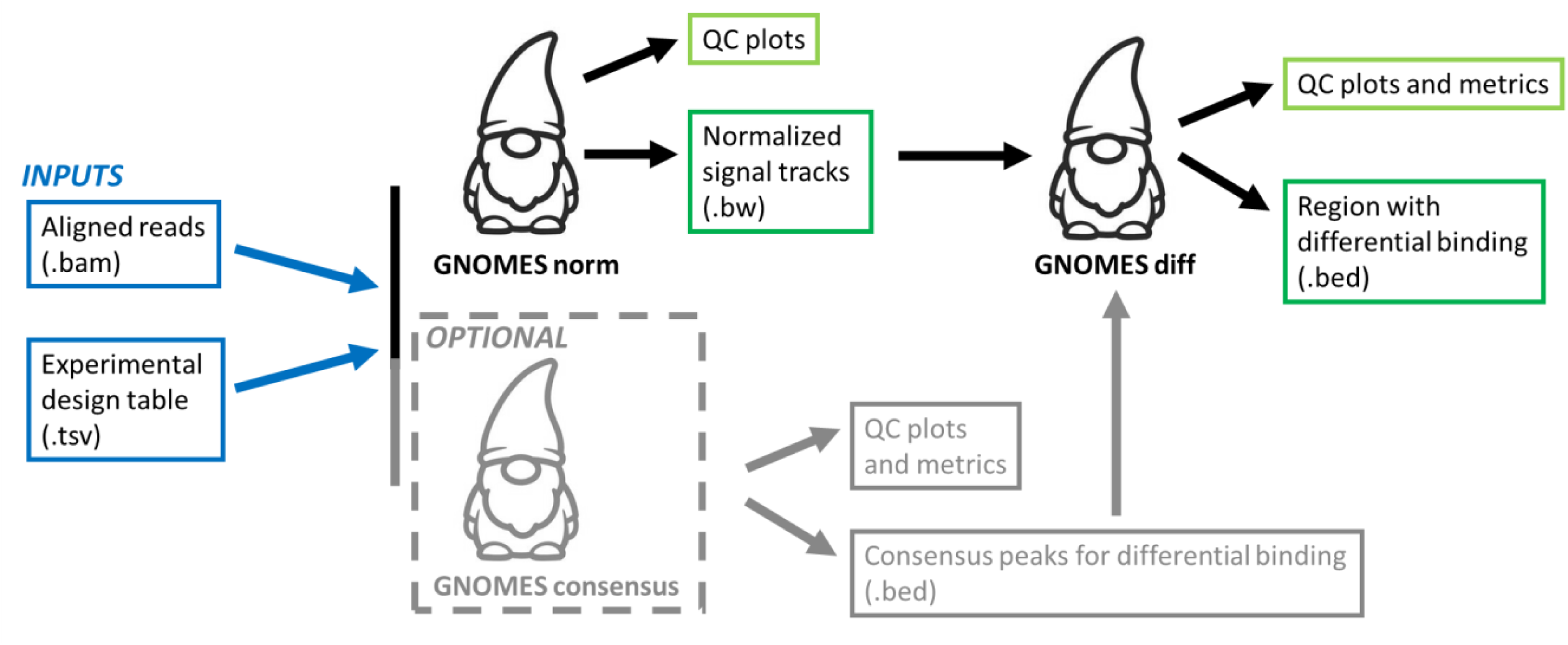
Overview of the *GNOMES* workflow. *GNOMES* is organized into three main modules: *norm*, *consensus*, and *diff*. The *norm* module performs signal normalization from aligned BAM files and generates normalized bigWig signal tracks and quality-control plots. The optional consensus module identifies candidate genomic regions that can be used as input for downstream differential binding analysis. Finally, the *diff* module performs differential binding testing within the selected regions using DESeq2 or edgeR and produces statistical result tables, diagnostic plots, and BED files describing regions showing gain or loss of signal.

### Input and preprocessing

*GNOMES* operates directly from aligned sequencing reads in BAM format, allowing users to integrate the framework easily into their existing ChIP-seq or CUT&RUN preprocessing pipelines. We recommend using high-quality uniquely aligned reads (ie. reads should be filtered to exclude unmapped reads, secondary alignments, low mapping quality reads, and reads failing quality checks). For example, commonly adopted tools such as Bowtie2 for alignment, samtools for read sorting and filtering, and Picard for duplicate removal can be used prior running *GNOMES* [14, 14, 15].

For each *GNOMES* module, the experimental design must be provided in a tab-separated metadata file describing the samples and associates BAM files (**Table1**). The metada includes information such as sample names, BAM file paths, experimental condition, and the chromatin target. Users can provide multiple chromatin target, and during the *norm* and *consensus* steps, *GNOMES* will automatically process samples independently for each target, preventing inappropriate comparisons between different chromatin marks. For the *diff* module, the same metadata file can be used, but the target should be specified using the --target flag. The target corresponds to the antibody used in the experiment (e.g., the histone mark or transcription factor immunoprecipitated in CUT&RUN or ChIP-seq). Control datasets (e.g., IGG or input) can be provided in the metadata file. In such case, control signal will be subtracted from the immunoprecipitation signal during the normalization step, and during the consensus step, controls will be used as background during MACS2 peak calling. Although *GNOMES* is not restricted to any specific organism, users must provide a tab-separated files containing chromosome size for their genome (**Table2**). Users can also provide a blacklist of genomic region in BED format to exclude them from the analysis.

**Table 1.** Example of GNOMES metadata file (--meta) provided as a tab-separated file.

| sample_id | bam | condition | target bam_control |
| --- | --- | --- | --- |
| WT_H3K27me3_1 | /path/WT_H3K27me3_1.bam | WT | /path/WT_input_1.bam |
| WT_H3K27me3_2 | /path/WT_H3K27me3_2.bam | WT | /path/WT_input_2.bam |
| KO_H3K27me3_1 | /path/KO_H3K27me3_1.bam | KO | /path/KO_input_1.bam |
| KO_H3K27me3_2 | /path/KO_H3K27me3_2.bam | KO | /path/KO_input_2.bam |
| WT_EZH2_1 | /path/WT_EZH2_1.bam | WT | /path/WT_input_1.bam |
| WT_EZH2_2 | /path/WT_EZH2_2.bam | WT | /path/WT_input_2.bam |
| KO_EZH2_1 | /path/KO_EZH2_1.bam | KO | /path/KO_input_1.bam |
| KO_EZH2_2 | /path/KO_EZH2_2.bam | KO | /path/KO_input_2.bam |

**Table 2.**
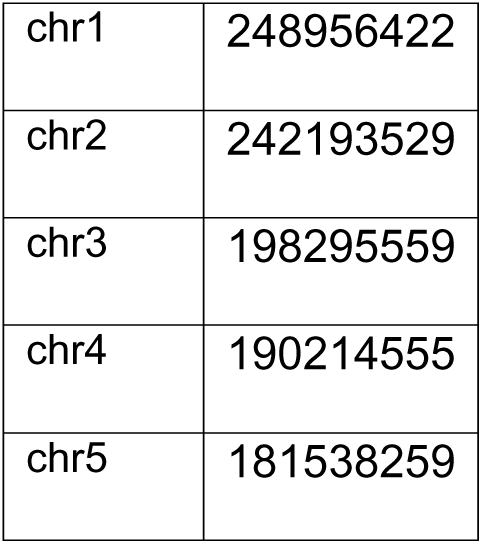
Example of chromosome size file (--chrom-sizes) containing chromosome names and lengths in base pairs provided as a tab-separated file.

|  |  |
| --- | --- |
| chr1 | 248956422 |
| chr2 | 242193529 |
| chr3 | 198295559 |
| chr4 | 190214555 |
| chr5 | 181538259 |

### Signal normalization

*GNOMES* applies a percentile-based normalization strategy to stabilize signal distributions across samples without assuming equal total chromatin occupancy between conditions [11] (**Figure2**). The normalization is implemented in the *GNOMES norm* module and operate directly from aligned BAM files. Briefly, high-quality aligned reads are first converted into BigWig format at 1bp resolution using bamCoverage. If a control dataset (e.g., IGG, input) is provided in the metada file, *GNOMES* uses bamCompare to subtract control signal from the corresponding immunoprecipitation (IP) sample [13]. When a blacklist of genomic regions is provided, overlapping signal is removed prior the normalization with bedtools intersect [16]. The resulting raw BigWig files are then converted to bedGraph format using bigWigToBedGraph [17]. From these files, *GNOMES* identifies all local maxima using a custom Python script. A local maximum is defined as a genomic bin with its signal value greater than that of the immediate upstream and downstream bins. *GNOMES* then computes the 99^th^ percentile (P99) of local maximum signal intensity for each sample. This value is used as a robust estimate of the upper signal distribution, while excluding the influence of extreme outliers from the 100^th^ percentile. By default, intermediate files such as local maxima and bedGraph files are removed after normalization to reduce storage usage, but they can be retained using –keep-temp. For a given target, *GNOMES* selects a reference sample (first sample from the metada file) and compute a scaling factor for each sample as:

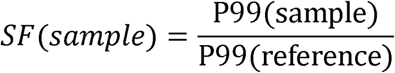

**Figure 2.**
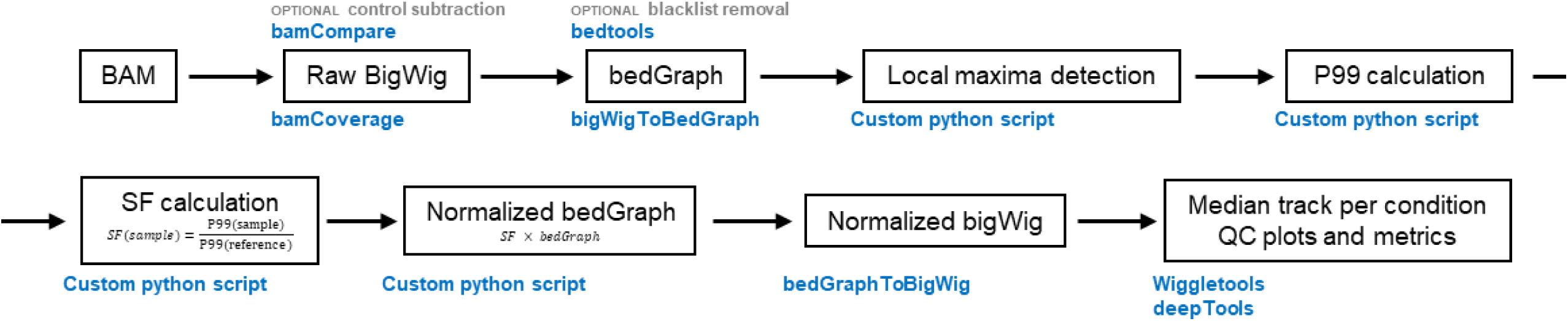
*GNOMES* signal normalization workflow. Aligned reads in BAM format are first converted into raw signal tracks in bigWig format using bamCoverage, or optionally bamCompare when control datasets (e.g., Input or IgG) are provided to subtract background signal. Raw bigWig files are converted to bedGraph format (bigWigToBedGraph), and optional blacklist filtering can be applied using bedtools. Local signal maxima are then identified using a custom Python script, and the 99th percentile (P99) of the local maxima distribution is calculated for each sample. These values are used to compute scaling factors for each sample. Scaling factors are applied to the bedGraph signal tracks to generate normalized bedGraph files, which are subsequently converted back to normalized bigWig tracks. Finally, *GNOMES* produces median signal tracks per condition using wiggletools, together with quality-control plots and metrics generated using deepTools to assess the effects of normalization across samples.

Bedgraph signal values are then multiplied by the corresponding sample scaling factor to generate normalized bedGraph tracks, wich are then converted back to normalized BigWig using bedgraphtoBigWig [17]. *GNOMES* also automatically generates median BigWig tracks for each condition and target using wiggletools [18]. Finally, the *norm* module produces quality-control metrics and diagnostic plots, including the scaling factors applied to each sample, as well as PCA and correlation heatmaps derived from both raw and normalized BigWig signals, generated using deepTools; allowing users to assess data quality before and after *GNOMES* normalization (**Figure2**).

### Consensus peak detection

After signal normalization, *GNOMES* includes an optional consensus module to generate candidate regions that will serve as input for the downstream binding analysis (**Figure3**). Consensus peaks correspond to genomic regions showing enrichment in at least one of the experimental conditions to be compared. As the genomic regions used for differential binding analysis directly influence the final set of differentially bound loci, the consensus module is designed to generate and compare multiple candidate peak sets. This flexibility is important because optimal peak-calling parameters often vary depending on dataset-specific features such as signal quality, complexity, and background noise [5].

**Figure 3.**
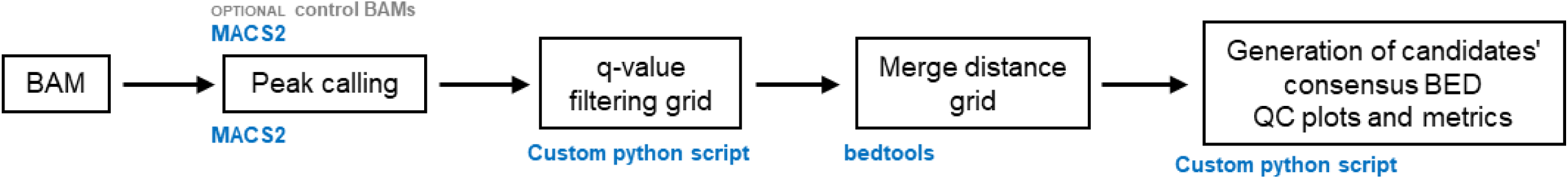
*GNOMES* consensus peak generation workflow. Aligned reads in BAM format are used for peak calling with MACS2, optionally with control BAM files (e.g., Input or IgG) as background during peak detection. Peaks identified across samples and conditions are subsequently filtered using a grid of user-defined q-value thresholds implemented through a custom Python script. The resulting peak sets are then merged across samples using bedtools with a range of region-merging distances to generate multiple candidate consensus peak sets. These steps produce a collection of BED files representing sets of genomic regions enriched in one or more conditions. *GNOMES* also generates summary metrics and quality-control plots to help selection of the most appropriate consensus region set for downstream differential binding analysis.

The *consensus* module performs peak calling with MACS2 using the same metadata table as the *norm* module. Samples from the same condition are pooled together in one peak calling step and represent a condition-level peak set. When control datasets are provided in the metadata file (e.g., IGG, input), they are supplied as MACS2 background during peak calling. All MACS2 parameters can be configured directly within the *GNOMES* framework, including the selection of narrow and broad peak detection modes. Once peaks are identified for each condition, they are concatenated, sorted, and merged using bedtools [16]; such that overlapping or nearby peaks can be combined into a single candidate region (**Figure3**). In practice, more stringent q-value thresholds reduce false-positive regions arising from background noise, whereas more permissive thresholds retain a larger number of candidate peaks. Similarly, the merge distance parameter controls whether nearby enriched regions are kept separate or combined into broader intervals. By default, five q-value thresholds (0.1, 0.05, 0.01, 0.001, and 0.0001) and five merge distances (0, 50, 100, 250, and 500 bp) are evaluated, producing 25 candidate consensus peak sets that users can compare to determine the most appropriate regions for downstream differential binding analysis.

In addition to the BED files themselves, the consensus module produces summary statistics for each candidate peak set, including the total number of peaks and peak width distributions. *GNOMES* also generates multi-page diagnostic plots showing the distribution of peak widths for each parameter tested. All parameters of the consensus workflow, including MACS2 mode, q-value thresholds, merge distances, and other peak-calling settings, are fully customizable, allowing the procedure to be adapted to a wide range of epigenomic datasets (**Figure3**).

### Differential binding analysis

Once normalized signal tracks have been generated, and an appropriate set of genomic regions selected, *GNOMES* performs differential binding analysis using its *diff* module (**Figure4**). This module requires the metadata table describing the experimental design, the chromatin target to be analyzed, a contrast specifying the conditions to compare (e.g., wild-type versus mutant), the directory containing normalized BigWig files, and a BED file defining the genomic regions in which differential binding will be tested. These regions may correspond to consensus peaks generated by the *GNOMES* consensus module, or to any user-provided set of genomic intervals. Consensus peak generation can also be performed directly within the *diff* workflow using MACS2 peak calling with user-defined parameters (including peak-calling mode, significance threshold, and merge distance). *GNOMES* uses deepTools computeMatrix to quantify the signal within each genomic region [13]. For each sample, the signal overlapping each region is converted into a count-like matrix suitable for downstream analysis. *GNOMES* implements two statistical methods for differential binding analysis: DESeq2 and edgeR [8, 9]. Both methods are fully configurable within the *GNOMES diff* command, allowing users to adjust significance thresholds, fold-change cutoffs, and low-count filtering parameters. Scaling procedures internal to the statistical models can also be customized (**Figure4**). For DESeq2, users may either allow standard estimation of size factors or disable it. Similarly, in edgeR users can choose among several scaling methods, including TMM, TMMwsp, RLE, or upperquartile, or disable scaling entirely (none). Allowing this step to be disabled is particularly useful in cases where large and unidirectional global changes in chromatin binding are expected (e.g., near-complete signal loss), as the assumption of balanced gains and losses may not hold for all epigenomic datasets [5].

**Figure 4.**
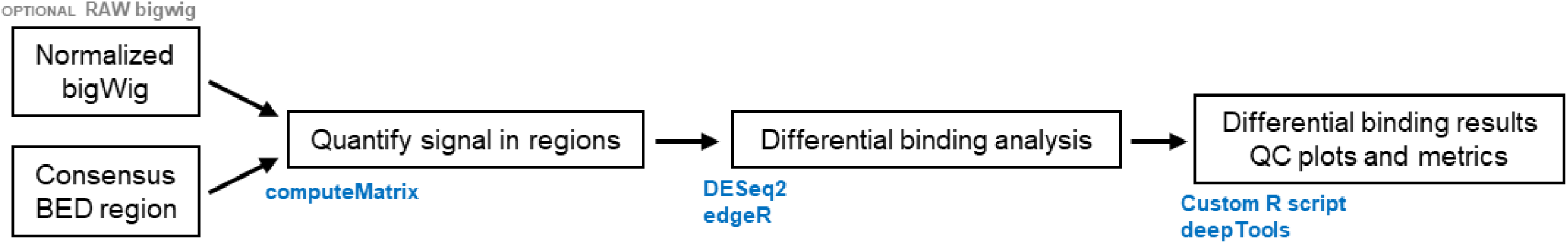
*GNOMES* differential binding analysis workflow. Normalized signal tracks in bigWig format are quantified across genomic regions from a BED file using deepTools computeMatrix. These regions may correspond to consensus peaks generated by the *GNOMES consensus* module or to any other user-provided genomic intervals. The resulting signal matrix is used for differential binding analysis using either DESeq2 or edgeR, implemented through custom R scripts. *GNOMES* then generates tables of differentially bound regions, together with BED files of regions showing gain or loss of signal between conditions. Additional quality-control and visualization outputs are produced, including diagnostic plots and heatmaps generated using deepTools, allowing users to assess the robustness and biological relevance of the differential binding results.

The *diff* module outputs both statistical result tables and a set of quality-control and result visualization plots. Differentially bound regions are reported with genomic coordinates, fold-change, and adjusted p-value; and are additionally exported as BED files of regions showing significant gain or loss of signal to facilitate rapid inspection in genome browser. Diagnostic plots include volcano plots, MA plots, PCA plots, and sample correlation heatmaps, enabling users to assess sample homogeneity, and relevance of significance thresholds chosen. Optionally, *GNOMES* also generates deepTools heatmaps and signal profile plots for significantly gained and lost regions, allowing direct assessment of global enrichment changes across samples. Because these visualization steps can be computationally intensive, they are implemented as optional outputs (**Figure4**).

Overall, the *diff* module was designed to provide an accessible yet highly customizable framework for differential binding analysis. By combining multiple statistical options with extensive quality-control outputs and visualization tools, *GNOMES* enables users to explore alternative parameter settings and select the analysis strategy that best matches the biological and technical characteristics of their dataset (**Figure4**).

## Results

### Identification of differential H3K27me3 regions during cerebellum development using GNOMES

To demonstrate *GNOMES* capabilities, we applied the workflow to publicly available H3K27me3 ChIP-seq data from mouse cerebellum at postnatal day 12 (P12) and postnatal day 21 (P21) [12]; two developmental stages characterized by substantial chromatin remodeling notably in the context of neuronal migration and synaptogenesis [19]. The dataset is single-end ChIP-seq data for the repressive histone mark H3K27me3 [20], and includes four biological replicates for P12 and five for P21. Matching bulk RNA-seq data are also available from the same study, and were later used to assess whether *GNOMES*-identified chromatin changes were associated with gene expression changes. First, we applied the *GNOMES norm* module to normalize the ChIP-seq signal using input samples as controls; when a control dataset (e.g., IgG or input) is specified in the metadata file, GNOMES performs control subtraction using bamCompare to removing background signal from the corresponding immunoprecipitation (IP) sample. Principal component analysis (PCA) of genome-wide signal tracks showed improved clustering of biological replicates by developmental stages after GNOMES normalization (**Figure5A**). In agreement, IGV snapshot of Hox gene clusters showed increased homogeneity of signal intensity across replicates after normalization (**Figure5B**). Next, we used *GNOMES consensus* module to identify candidate genomic regions for differential binding analysis. As H3K27me3 typically forms broad domains [20], peaks were called using MACS2 broad peak mode. We tested combinations of five q-value and merge distance thresholds (--macs2-qvalue 0.01, 0.001, 0.0001, 0.00001, 0.000001; --macs2-merge 0, 50, 100, 250, 500). *GNOMES* automatically summarized the resulting peak sets in a grid of candidate consensus regions (**Supplementary Table1**). From manual inspection on IGV genome browser, we decided to use a q-value threshold of 1e-5 with a 100bp peak merge distance, as it provides the best balance between retaining true peaks while removing the noisy ones (**Figure5C**). The selected consensus set contained 11,107 candidate regions with a median size of approximately 3kb (**Supplementary Figure1A**). Finally, differential binding analysis was performed with the *GNOMES diff* module using edgeR and upperquartile scaling (--diff-method edger; --edger-min-counts 100; --edger-norm upperquartile). This analysis identified 2,174 differentially bound regions between P12 and P21 using an adjusted *p*-value threshold of 0.05 and an absolute log2 fold-change cutoff of 0.58 (--edger-alpha 0.05; --edger-lfc 0.58), including 679 regions with decreased enrichment and 1,495 regions with increased enrichment from P12 to P21 (**Figure5D; Supplementary Figure1B**). Inspection of these significant regions in genome browser confirms clear changes in chromatin signal across biological conditions, consistent with the patterns observed in the signal heatmaps (**Figure 5E**). Furthermore, correlation analysis of counts within consensus peak regions showed improved condition-based clustering of sample when using *GNOMES*-normalized counts as compared with raw counts (**Supplementary Figure1C**).

**Figure 5.**
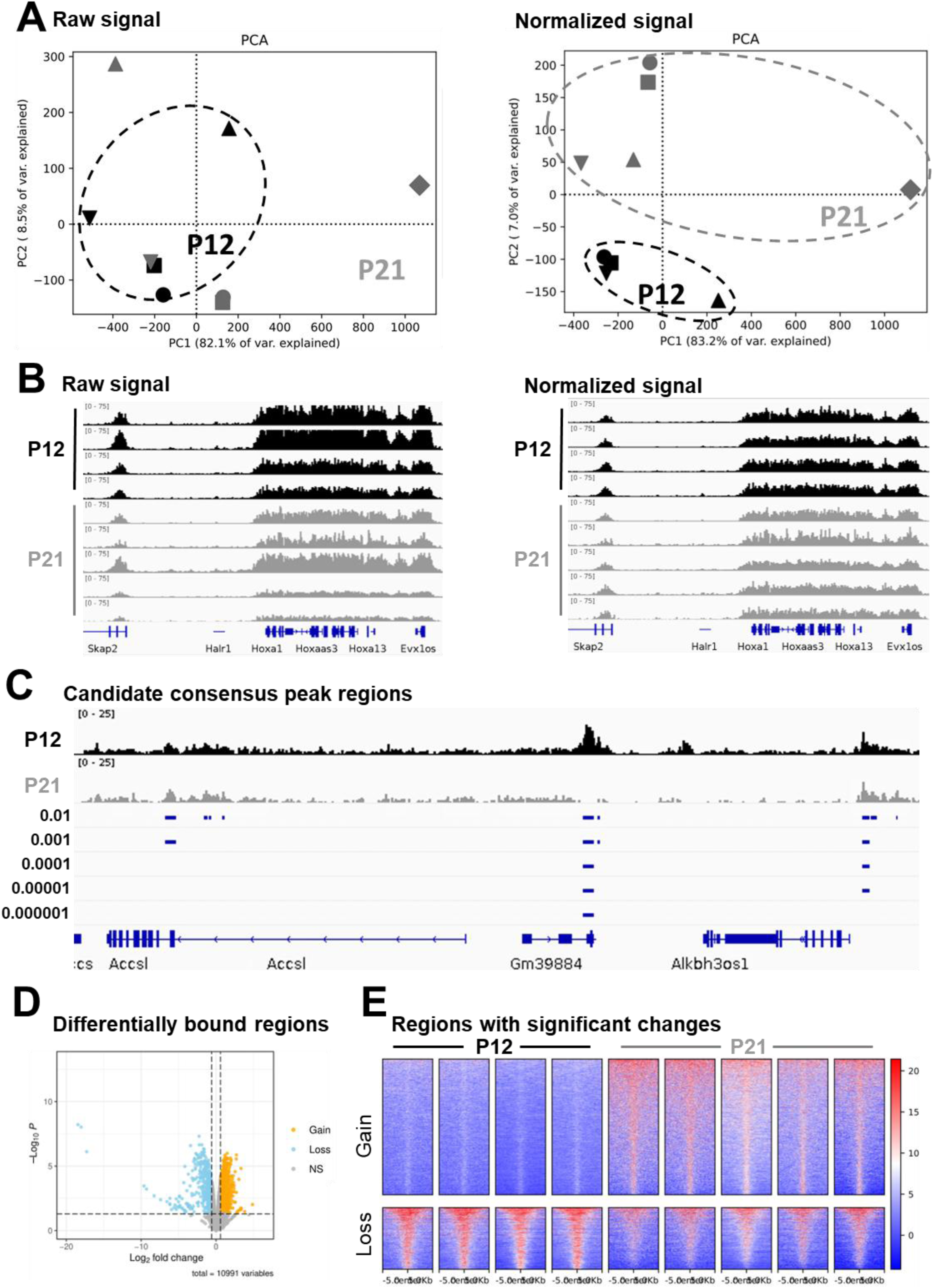
GNOMES normalization and differential H3K27me3 binding analysis between P12 and P21 mouse cerebellum. **(A)** Principal Component Analysis (PCA) of genome-wide H3K27me3 signal derived from raw (left) and normalized (right) BigWig tracks. **(B)** IGV snapshot of Hox gene loci before (left) and after (right) normalization. **(C)** IGV snapshot of candidate consensus peak regions generated by the *GNOMES consensus* module using multiple MACS2 q-value thresholds (0.01-0.000001) with a 100bp merge distance. **(D)** Volcano plot of H3K27me3 differentially bound regions from P12 to P21. Significantly gain and loss regions are highlighted in orange and blue, respectively (edgeR, *adjusted p*-value < 0.05 and |log_₂_FC| ≥ 0.58). **(E)** Heatmap of normalized H3K27me3 signal centered on consensus peaks with significant H3K27me3 binding changes from P12 to P21.

Together, these results demonstrate that *GNOMES* integrate an easy to use, and highly customizable workflow, for epigenomic signal normalization and differential binding; producing key summary metrics and publication-ready figures.

### GNOMES-identified H3K27me3 changes are associated with transcriptional regulation of neurodevelopmental genes

To assess whether the H3K27me3 binding changes identified by *GNOMES* are relevant, we examined whether these chromatin changes were associated with gene expression changes. Indeed, H3K27me3 being a well-established repressive histone mark, we expected genes gaining H3K27me3 to show decreased expression, whereas gene losing H3K27me3 would show increased expression [20].

To test this, we retrieved and re-analyzed the bulk RNA-seq data generated from the same mouse cerebellum study at P12 and P21 [12]. Using DEseq2 with an adjusted *p*-value threshold of 0.05 and an absolute log2 fold-change cutoff of 0.58, we identified 5,138 differentially expressed genes, including 2,919 down- and 2,219 up- regulated genes from P12 to P21 (**Supplementary Figure2A and 2B**). Then, we examined with ChIPseeker the genomic distribution of *GNOMES* H3K27me3 differentially bound regions [21]. Interestingly, regions losing H3K27me3 were strongly enriched at promoters, whereas regions gaining H3K27me3 were more frequently associated with intergenic regions (**Figure6A**). Because chromatin changes at promoters are most likely to directly influence transcription, we focused on genes whose promoter or 5′ regulatory regions overlapped *GNOMES*-identified differential H3K27me3 regions. Among the 1,495 regions gaining H3K27me3, 229 unique genes showed promoter-proximal enrichment, whereas 482 unique genes were associated with the 679 regions losing H3K27me3. Aggregated H3K27me3 signal on these genes confirmed clear changes in H3K27me3 enrichment from P12 to P21 (**Figure6B**). We next examined the gene expression changes occurring from P12 to P21 on these same genes. Remarkably, genes gaining H3K27me3 in their proximal-promoter regions were predominantly significantly down-regulated (126/229), whereas genes losing H3K27me3 were mostly significantly up-regulated (154/482) (**Figure6C**). The effect was more pronounced for genes gaining H3K27me3, consistent with the strong chromatin compaction associated with this mark, which can hinder transcription factor (TF) binding [22]. In contrast, loss of H3K27me3 alone does not necessarily activate transcription, as gene activation requires accessible chromatin and recruitment of activating TFs [23].

To further evaluate the biological relevance of these gene sets, we performed Gene Ontology (GO) enrichment analysis with enrichR [24]. Down-regulated genes gaining H3K27me3 were significantly enriched for developmental processes including Regulation of neurogenesis, gliogenesis, and embryonic organ morphogenesis (**Figure 6D**). Conversely, up-regulated genes losing H3K27me3 were enriched for signaling pathways including small GTPase mediated signal transduction, Ras protein signal transduction, and canonical Wnt signaling pathway (**Figure 6D**); processes known to be involved in neuronal maturation and signaling during cerebellar development [25, 26]. Visualization of representative loci in a genome browser further confirmed the relationship between H3K27me3 changes and gene expression. For example, genes such as Sox9, Mycn, and Sox11, which show decreased expression at P21, display clear increases in H3K27me3 signal across their loci (**Figure6E**). In contrast, genes including Wnt7a, Bdnf, and Nr4a2, which become transcriptionally activated during cerebellum maturation, show marked reductions in H3K27me3 signal at P21.

**Figure 6.**
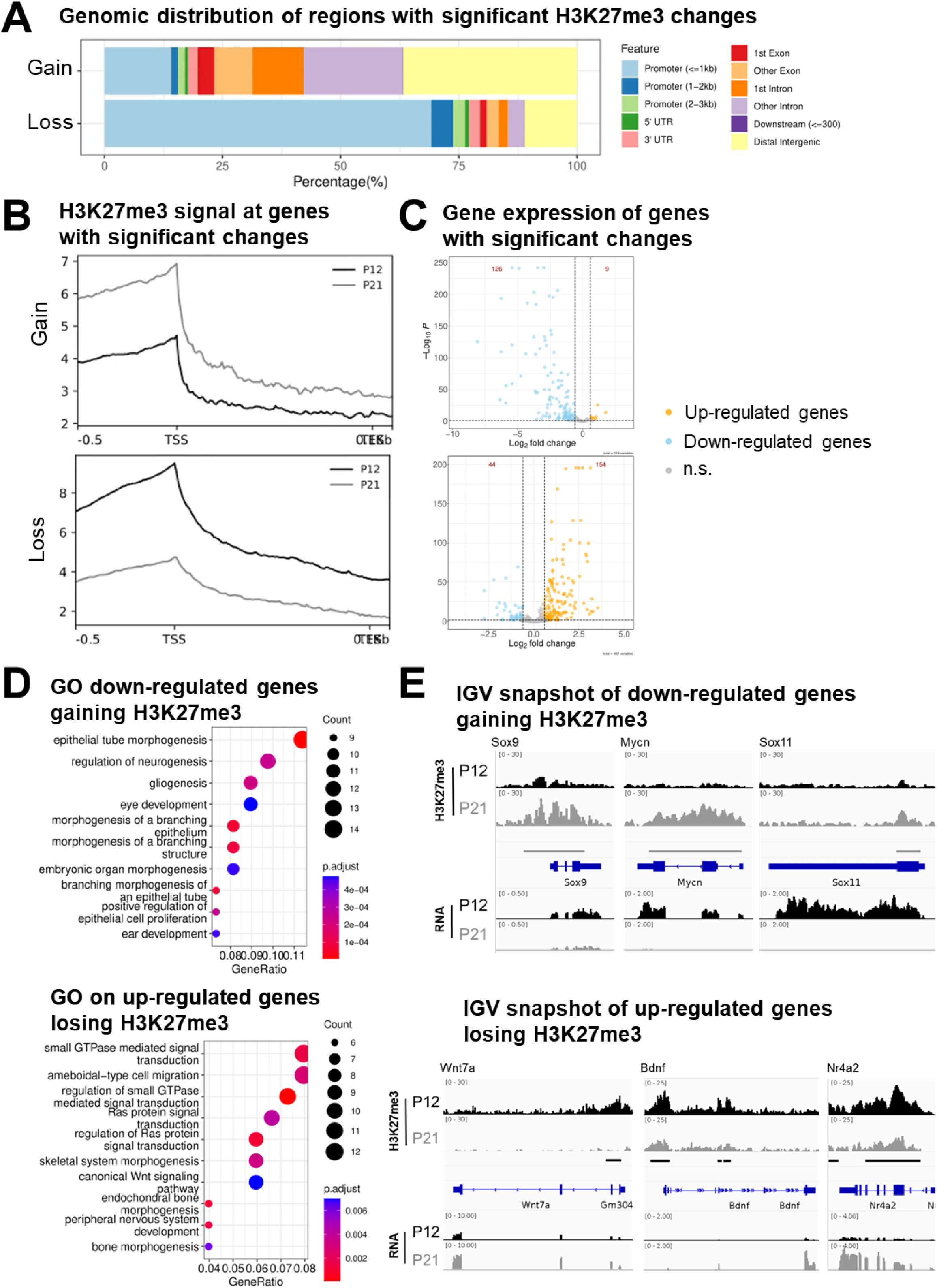
H3K27me3-mediated changes of gene expression between P12 and P21 mouse cerebellum. **(A)** Genomic distribution of regions with significant H3K27me3 binding changes from P12 to P21. Peak were annotated relative to genomic features using ChIPseeker. **(B)** Average H3K27me3 signal across genes containing significant H3K27me3 binding changes within their promoter and/or 5’ regions. **(C)** Volcano plot expression of genes with significant H3K27me3 binding changes displayed in B). Significantly up-regulated and down-regulated genes are highlighted in orange and blue, respectively (*adjusted p*-value < 0.05 and |log_₂_FC| ≥ 0.58), with the total number of DEGs indicated in red. **(D)** Gene Ontology (GO) biological process enrichment analysis on down-regulated genes gaining H3K27me3 (top) and up-regulated genes losing H3K27me3 (bottom) from P12 to P21. **(E)** IGV snapshot showing H3K27me3 signal and RNA abundance of genes down-regulated and gaining H3K27me3 (top), and genes up-regulated and losing H3K27me3 (bottom) from P12 to P21.

Together, these results demonstrate that *GNOMES* differential H3K27me3 regions correspond to biologically meaningful chromatin changes associated with transcriptional regulation during cerebellum development. The strong correlation between chromatin state changes and gene expression dynamics provides independent validation that *GNOMES* accurately identifies functional epigenomic alterations.

## Discussion

### *GNOMES* provides a flexible and accessible framework for epigenomic normalization and differential binding analysis

*GNOMES* provides an accessible, robust, and highly customizable workflow for epigenomic signal normalization and differential binding analysis. The framework integrates multiple widely used tools, including MACS2, deepTools, DESeq2, and edgeR, within a unified pipeline implemented in Python and R [6, 8, 9, 13]. The complete workflow can be executed from only three command-line steps, including one optional. The software supports installation through Conda environments and Apptainer containers, allowing easy installation on any computational environment. Furthermore, *GNOMES* is open source and maintained through a public GitHub repository, where users can report issues, contribute improvements and follow development.

A key feature of *GNOMES* is its normalization strategy based on local maxima epigenomic signal, which rescale samples to have comparable signal ranges, ensuring different samples remain within the same order of magnitude. This facilitates meaningful comparisons across conditions, while maintaining true biological changes, even in the context of near complete loss or gain of signals. Furthermore, *GNOMES* produces both normalized signal tracks and differential binding results allowing the user to directly verify differential regions within genome browsers.

Although the use of input controls is recommended, *GNOMES* can also operate without control samples, increasing its flexibility for datasets where input libraries are unavailable. The framework has been tested on both ChIP-seq and CUT&RUN datasets, where it produces robust normalization and consistent differential binding results. *GNOMES* is organism-agnostic and requires only uniquely aligned reads as input, allowing it to be applied to a wide range of species and experimental systems. In addition, the pipeline automatically generates multiple quality-control metrics and publication-ready plots that facilitate interpretation of the results.

### Limitations and future directions

Despite these advantages, several limitations and potential areas for improvement remain. First, while *GNOMES* has been validated on ChIP-seq and CUT&RUN datasets, it has not been tested on CUT&TAG. Future development could also expand the functionality of the consensus peak module. Currently, *GNOMES* relies on the widely used MACS2 peak caller to identify candidate regions [6]. Although MACS2 already provides extensive parameter flexibility, integrating additional peak-calling algorithms such as SEACR and HOMER [27, 28], could further broaden the applicability of the workflow to different experimental designs or signal profiles.

Another potential extension would be to include automated annotation of peaks to nearby genes. However, several well-established tools, such as ChIPseeker [21], already perform this task effectively. Integrating such functionality directly into *GNOMES* could increase convenience but might also reduce the organism-agnostic nature of the framework, which currently represents one of its strengths.

Finally, as with any computational analysis pipeline: *GNOMES* relies on the quality of the input data. While the normalization strategy improves comparability between samples, it cannot compensate for severely compromised or low-quality sequencing data. Careful experimental design and rigorous data quality control therefore remain essential for reliable epigenomic analysis.

## Conclusion

*GNOMES* provides an integrated framework for normalization, visualization, and differential binding analysis of epigenomic datasets. By combining robust signal normalization with peak-centric statistical testing and extensive quality-control outputs, *GNOMES* facilitates reproducible and interpretable analysis of ChIP-seq and CUT&RUN experiments.

## Availability and requirements

**Project name**: GNOMES (Genome-wide NOrmalization of Mapped Epigenomic Signals)

**Project home page**: https://github.com/RouleThomas/GNOMES

**Operating system(s)**: Linux, macOS (Unix-based systems recommended)

**Programming language**: Bash, Python, R

**Other requirements**: Python (≥3.8), deepTools, bedtools, wiggletools, UCSC utilities (bigWigToBedGraph, bedGraphToBigWig), MACS2, Conda

**License**: MIT License

**Any restrictions to use by non-academics**: None (freely available for academic and non-academic use)

## Authors’ Contributions

T.R.: Developed the software, performed the analyses, and drafted the manuscript. N.A.: Supervised the study and contributed to methodological refinement and manuscript writing (review and editing). All authors approved the final manuscript.

## Acknowledgments

We thank Jasmine Akoto for testing the tool and providing valuable feedback.

## Data availability

The *GNOMES* source code is available on GitHub at https://github.com/RouleThomas/GNOMES. Commands used to reproduce the *GNOMES* analysis presented in the Results section are provided in the example_workflow/ directory of the *GNOMES* GitHub repository.

## Figure legends

**Supplementary Figure 1. GNOMES consensus and diff modules quality control plots.**

**(A)** Peak width distribution of the consensus peak set selected (MACS2 q-value threshold of 0.00001 and a merge distance of 100bp) for downstream differential binding analysis. (B) MA plot of differentially H3K27me3 bound regions from P12 to P21. Significantly gain and loss regions are highlighted in orange and blue, respectively (edgeR, *adjusted p*-value < 0.05 and |log_₂_FC| ≥ 0.58). (C) Sample correlation heatmap of signal in consensus peaks before (left) and after (right) *GNOMES* normalization.

**Supplementary Figure 2. Bulk-RNAseq differential expression analysis in P12 and P21 mouse cerebellum samples.**

**(A**) Principal Component Analysis (PCA) of bulk-RNAseq samples after variance-stabilizing transformation with DESeq2. **(B)** Volcano plot of differentially expressed genes (DEGs) between P12 and P21. Significantly up-regulated and down-regulated genes are highlighted in orange and blue, respectively (*adjusted p*-value < 0.05 and |log_₂_FC| ≥ 0.58), with the total number of DEGs indicated in red.

## References

1. Lloyd SM, Bao X. Pinpointing the Genomic Localizations of Chromatin-Associated Proteins: The Yesterday, Today, and Tomorrow of ChIP_-_seq. CP Cell Biology. 2019;84:e89. 10.1002/cpcb.89.

2. Park PJ. ChIP–seq: advantages and challenges of a maturing technology. Nat Rev Genet. 2009;10:669–80. 10.1038/nrg2641.

3. Allhoff M, Seré K, F. Pires J, Zenke M, G. Costa I. Differential peak calling of ChIP-seq signals with replicates with THOR. Nucleic Acids Res. 2016;:gkw680. 10.1093/nar/gkw680.

4. Lun ATL, Smyth GK. csaw: a Bioconductor package for differential binding analysis of ChIP-seq data using sliding windows. Nucleic Acids Research. 2016;44:e45–e45. 10.1093/nar/gkv1191.

5. Colando S, Schulz D, Hardin J. Selecting ChIP-seq normalization methods from the perspective of their technical conditions. Briefings in Bioinformatics. 2025;26:bbaf431. 10.1093/bib/bbaf431.

6. Zhang Y, Liu T, Meyer CA, Eeckhoute J, Johnson DS, Bernstein BE, et al. Model-based Analysis of ChIP-Seq (MACS). Genome Biol. 2008;9:R137. 10.1186/gb-2008-9-9-r137.

7. Zang C, Schones DE, Zeng C, Cui K, Zhao K, Peng W. A clustering approach for identification of enriched domains from histone modification ChIP-Seq data. Bioinformatics. 2009;25:1952–8. 10.1093/bioinformatics/btp340.

8. Love MI, Huber W, Anders S. Moderated estimation of fold change and dispersion for RNA-seq data with DESeq2. Genome Biol. 2014;15:550. 10.1186/s13059-014-0550-8.

9. Robinson MD, McCarthy DJ, Smyth GK. edgeR : a Bioconductor package for differential expression analysis of digital gene expression data. Bioinformatics. 2010;26:139–40. 10.1093/bioinformatics/btp616.

10. Stark R, Brown G. Di Bind : di erential binding analysis of ChIP-Seq peak data.

11. Parmar A, Srinivasan A, Krockenberger L, Augustine A, Gong O, Bullard AC, et al. Polycomb repressive complexes 1 and 2 independently and dynamically regulate euchromatin during cerebellar neurodevelopment. PLoS Genet. 2025;21:e1011843. 10.1371/journal.pgen.1011843.

12. Mätlik K, Govek E-E, Paul MR, Allis CD, Hatten ME. Histone bivalency regulates the timing of cerebellar granule cell development. Genes Dev. 2023;37:570–89. 10.1101/gad.350594.123.

13. Ramírez F, Ryan DP, Grüning B, Bhardwaj V, Kilpert F, Richter AS, et al. deepTools2: a next generation web server for deep-sequencing data analysis. Nucleic Acids Res. 2016;44:W160–5. 10.1093/nar/gkw257.

14. Langmead B, Salzberg SL. Fast gapped-read alignment with Bowtie 2. Nat Methods. 2012;9:357–9. 10.1038/nmeth.1923.

15. Li H, Handsaker B, Wysoker A, Fennell T, Ruan J, Homer N, et al. The Sequence Alignment/Map format and SAMtools. Bioinformatics. 2009;25:2078–9. 10.1093/bioinformatics/btp352.

16. Quinlan AR, Hall IM. BEDTools: a flexible suite of utilities for comparing genomic features. Bioinformatics. 2010;26:841–2. 10.1093/bioinformatics/btq033.

17. Kent WJ, Zweig AS, Barber G, Hinrichs AS, Karolchik D. BigWig and BigBed: enabling browsing of large distributed datasets. Bioinformatics. 2010;26:2204–7. 10.1093/bioinformatics/btq351.

18. Zerbino DR, Johnson N, Juettemann T, Wilder SP, Flicek P. WiggleTools: parallel processing of large collections of genome-wide datasets for visualization and statistical analysis. Bioinformatics. 2014;30:1008–9. 10.1093/bioinformatics/btt737.

19. Akoto J, Roule T, Akizu N. PRC2 Diversity in Neuronal Differentiation and Developmental Disorders. Genes. 2025;16:1191. 10.3390/genes16101191.

20. Wiles ET, Selker EU. H3K27 methylation: a promiscuous repressive chromatin mark. Current Opinion in Genetics & Development. 2017;43:31–7. 10.1016/j.gde.2016.11.001.

21. Yu G, Wang L-G, He Q-Y. ChIPseeker: an R/Bioconductor package for ChIP peak annotation, comparison and visualization. Bioinformatics. 2015;31:2382–3. 10.1093/bioinformatics/btv145.

22. Margueron R, Reinberg D. The Polycomb complex PRC2 and its mark in life. Nature. 2011;469:343–9. 10.1038/nature09784.

23. Bieluszewski T, Prakash S, Roulé T, Wagner D. The Role and Activity of SWI/SNF Chromatin Remodelers. Annu Rev Plant Biol. 2023;74:139–63. 10.1146/annurev-arplant-102820-093218.

24. Xie Z, Bailey A, Kuleshov MV, Clarke DJB, Evangelista JE, Jenkins SL, et al. Gene Set Knowledge Discovery with Enrichr. Current Protocols. 2021;1:e90. 10.1002/cpz1.90.

25. Hall A, Lalli G. Rho and Ras GTPases in Axon Growth, Guidance, and Branching. Cold Spring Harbor Perspectives in Biology. 2010;2:a001818–a001818. 10.1101/cshperspect.a001818.

26. Leto K, Arancillo M, Becker EBE, Buffo A, Chiang C, Ding B, et al. Consensus Paper: Cerebellar Development. Cerebellum. 2016;15:789–828. 10.1007/s12311-015-0724-2.

27. Meers MP, Tenenbaum D, Henikoff S. Peak calling by Sparse Enrichment Analysis for CUT&RUN chromatin profiling. Epigenetics & Chromatin. 2019;12:42. 10.1186/s13072-019-0287-4.

28. Heinz S, Benner C, Spann N, Bertolino E, Lin YC, Laslo P, et al. Simple Combinations of Lineage-Determining Transcription Factors Prime cis-Regulatory Elements Required for Macrophage and B Cell Identities. Molecular Cell. 2010;38:576–89. 10.1016/j.molcel.2010.05.004.

