## Supplementary Figure1 for "GNOMES: an integrated framework for genome-wide normalization and differential binding analysis of CUT&RUN and ChIP-seq data"

**Suppl. Figure 1**

**A**

**Consensus peak width distribution**

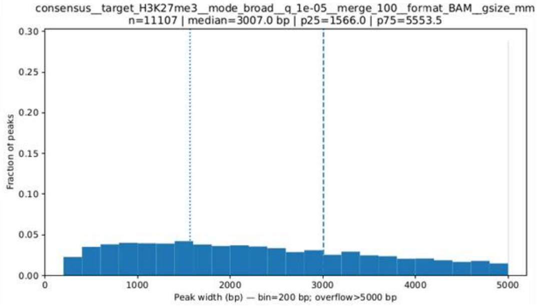


**B**

**MA plot of differentially bound regions**

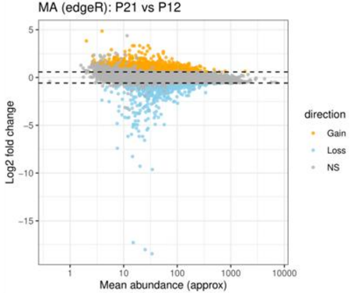


**C**

**Sample correlation heatmap**

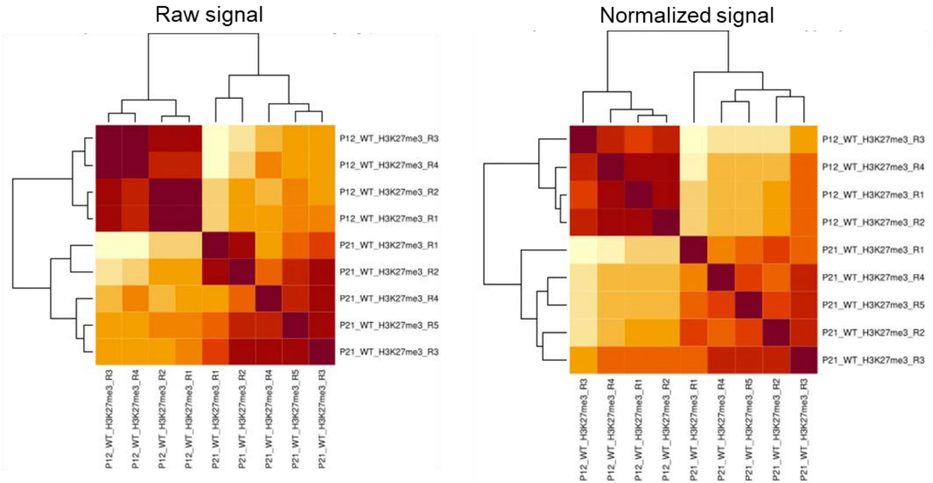
