## Supplementary figures and images for "GNOMES: an integrated framework for genome-wide normalization and differential binding analysis of CUT&RUN and ChIP-seq data"

### Supplementary Figure2

**Suppl. Figure 2**

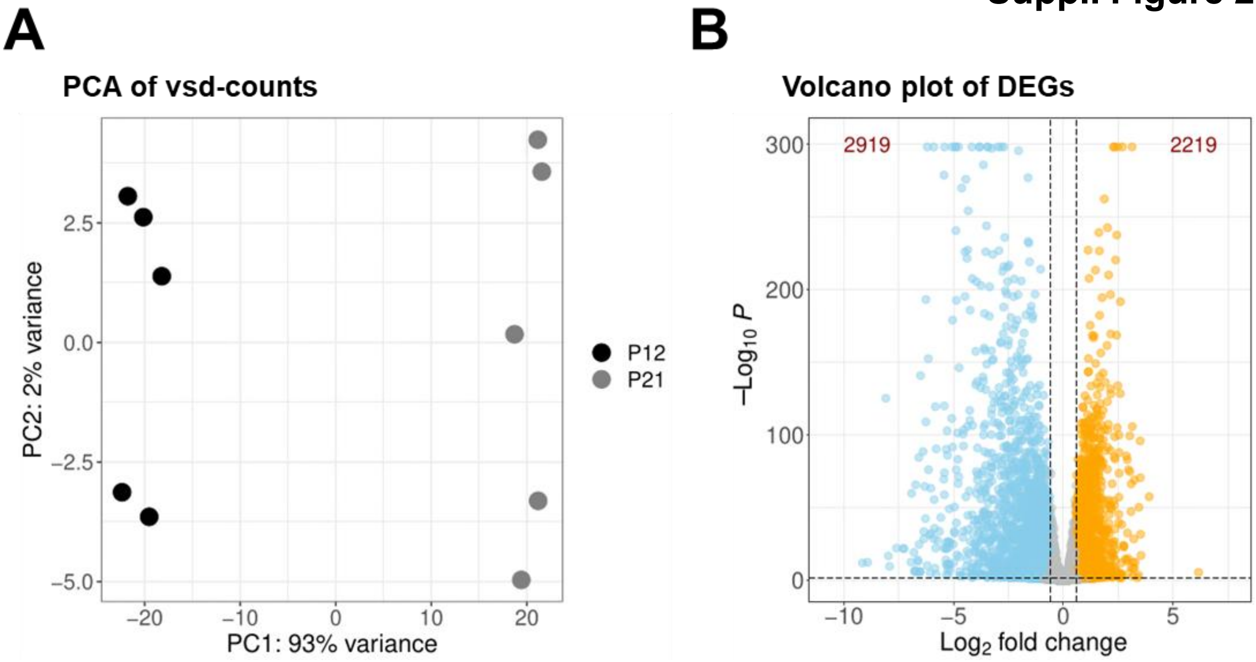
